# Assessing alternative methods of using population genomic data to measure changes in population size

**DOI:** 10.64898/2026.03.27.714834

**Authors:** Laiyin Zhou, Tin-Yu J. Hui, Austin Burt

**Affiliations:** Department of Life Sciences, Silwood Park Campus, Imperial College London, Ascot, United Kingdom

## Abstract

Malaria remains a major global health burden, with traditional control methods facing challenges such as insecticide resistance and high operational costs. Genetic biocontrol offers a promising alternative for mosquito population suppression, but its field efficacy would require assessment. This study evaluates the role that population genomic statistics can play in detecting decreases in population size in the context of a cluster randomized control trial (cRCT), investigating the response of nucleotide diversity (*π*), Tajima’s D, segregating sites, and linkage disequilibrium (LD) under both constant and seasonal demographic scenarios. We simulated 90% and 99% population declines with various degrees of between-cluster heterogeneity, and assessed the detection power of each statistic over time and number of clusters per arm. Results show that Tajima’s D is highly sensitive and robust across crash severity, seasonality and heterogeneity scenarios. Segregating sites has similar power to Tajima’s D when baseline data are available. We further estimated that cRCTs require approximately 3 to 5 villages per treatment arm to achieve adequate statistical power. These findings provide recommendations for genetic monitoring of vector control interventions in wild populations.

## 1 Introduction

Malaria is one of the most devastating global health challenges, responsible for approximately 600,000 deaths annually, 95% of which occurred in the African Region (World Health Organization, 2025). Despite decades of concerted efforts, progress in malaria control has stagnated since 2015 in part due to emerging insecticide resistance, antimalarial drug resistance, logistical challenges in sustaining conventional interventions, outdoor transmission, and funding gaps (Dhiman, 2019; Ohiri et al., 2022). Common control strategies, including drugs and vaccines, bed nets and insecticides, as well as regional infrastructure improvements, often require continuous financial and operational investments (Hemingway et al., 2016). Their effectiveness is further constrained by infrastructural limitations and diminishing returns in high-transmission settings (Cohen et al., 2022).

The limitations of existing approaches have spurred interest in novel vector control strategies such as genetic biocontrol. Early applications of genetic strategies for pest control include the use of genetically modified sterile insects. In parallel, gene drive systems targeting *Anopheles gambiae s.l.* mosquitoes, a species complex containing the major malaria vectors in Sub-Saharan Africa, have shown potentials in laboratory, though no gene drive mosquito has yet been released into the wild (Schairer et al., 2021). Gene drives leverage CRISPR-Cas9 technology to bias the inheritance of engineered traits for knocking out genes responsible for essential functions (e.g. sex-determining or fertility), or inserting transmission blocking effectors, potentially enabling rapid population-wide suppression or modification within a few generations (Tajudeen et al., 2023). As promising results have been shown in caged experiments and mathematical modelling, the current emphasis is on strain characterisation, assessment of efficacy and biosafety, and regulatory and community approval on future field trials. (Hammond et al., 2016; Hancock et al., 2024).

Clustered randomised control trials (cRCT) are the gold standard for evaluating the efficacy of new vector control interventions (Puffer et al., 2005). In this study design groups or clusters of individuals, rather than individuals, are randomly assigned to intervention or control conditions. This design is particularly valuable when interventions are naturally delivered at cluster level, or when individual randomisation is impractical or would risk contamination between study arms. Many existing cRCTs (or RCTs in general) have assessed the suppression of vector densities as one of their entomological endpoints (Belay et al., 2025; Staedke et al., 2020; Yovogan et al., 2023), or as the basis for sample size and power calculation (Hancock et al., 2025). Vector densities are typically estimated by conventional trapping methods (e.g. light traps and indoor spray catch), although Mark-Release-Recapture (MRR) and Close-Kin Mark Recapture (CKMR) methods also exist (Epopa et al., 2017; Sharma et al., 2022). These are measures of the census size *N_c_*, defined as the number of adult in the focal population (Luikart et al., 2010), which can be highly variable for mosquitoes. Seasonal dynamics of the underlying populations driven by fresh water availability, alongside other abiotic factors such as land use and climatic zones, and random noise from the trapping method, are the contributors to these variabilities (Hui et al., 2025). Previous estimates of *N_c_* suggested an order of magnitude difference between the rainy and dry season sizes in the Sahel region of West Africa (Epopa et al., 2017), and vector composition within *An. gambiae s.l.* can also vary across nearby villages *<*50km apart. These may potentially mask the intervention effect, resulting in low statistical power, or needing a large trial size.

Such suppression effects should also be reflected in the effective population size *N_e_*, a genetic measure linking up the magnitude of genetic drift of the focal population to the idealised one, which also governs the level of genetic diversity, rate of evolution, correlation of genetic loci etc. (Charlesworth, 2009). The additional investment in DNA sequencing enables population monitoring through genetic summary statistics, which capture complementary aspects of the genetic signal spatially and temporally, thereby offering the potential to reduce sampling noise. They were known to respond to *N_e_* in applications such as conservation and demographic estimation (Antao et al., 2011; Hui, 2025; Mwima et al., 2024; O’Loughlin et al., 2016), although their use in cRCT settings remains a relatively new concept.

To address this gap, this study aims to: (1) Identify which genetic summary statistics are useful for detecting mosquito population declines in the context of a cRCT, (2) Examine the effect of spatial and temporal heterogeneity in terms of seasonal and between-cluster variations, and (3) Recommend cRCT sample size requirements (number of clusters per treatment arm) under different scenarios.

## 2 Methods

### 2.1 Genome simulations

We used *msprime* (v 1.3.4) (Baumdicker et al., 2022) to simulate genomes for the clusters under various chosen scenarios (e.g. constant or seasonal model, and heterogeneity, see below). Each independent run simulates the genetic contents of a population (cluster) over time under a given scenario until t=36 generations ago. For those receiving the intervention, an immediate population crash was experienced from t=36 to t=0 according to the intended suppression level. Those in the control arm remained intact. We sampled 50 diploid individuals every 3 generations, from the present day (t=0) up to t=48 generations ago, covering both the pre- and post-intervention periods. Note that 1 year equates to about 10-12 mosquito generations (Hui et al., 2021). Two chromosomes of 1Mb each were simulated, and the mutation rate (*µ*) was set to 2.5 *×* 10*^−^*^8^ per base per generation, and recombination rate *r* = 10 *× µ* between adjacent base pairs (Pombi et al., 2006). To accurately capture recent genealogical processes while maintaining computation efficiency, we implemented a hybrid simulation approach: the discrete-time Wright-Fisher (DTWF) model (Nelson et al., 2020) was used for the most recent 50 generations, followed by the standard Hudson coalescent model for deeper coalescence. A total of 200 control and 200 intervention clusters were simulated under each model, from which *k* samples per arm were sampled to calculate the power of a cRCT at each sampling time post-crash (*t*_2_).

#### The constant population model

This model assumes a constant pre-intervention *N_e_* of 30,000. Those receiving the intervention will have their *N_e_* reduced to 3,000 for 90% suppression, or 300 for 99% suppression (Figure 1A).

**Figure 1:**
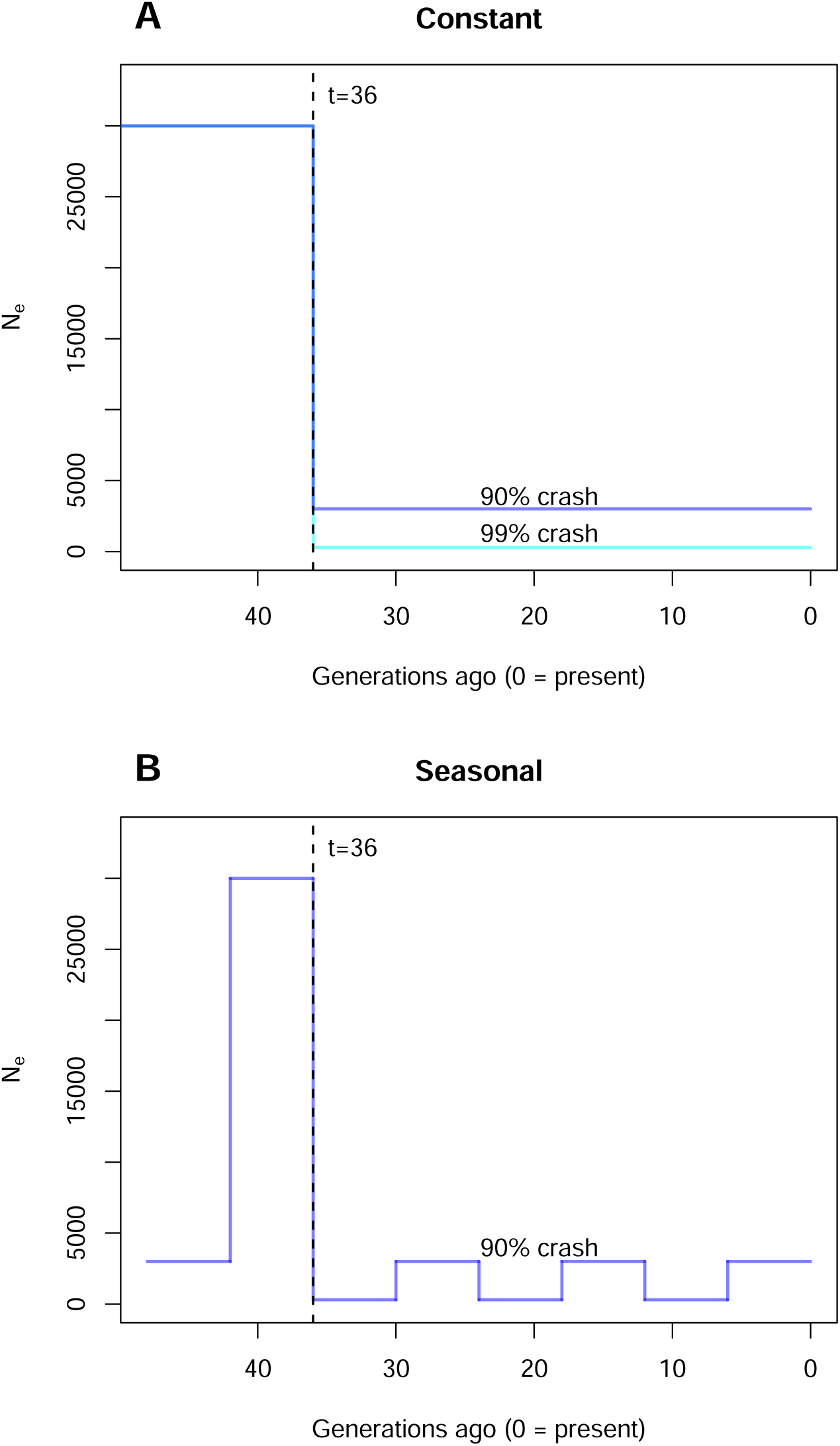
Schematic representation of demographic and intervention scenarios in (A) constant and (B) seasonal populations on *msprime*. Generations are counted backward in time, with t=0 being the most recent generation. Intervention is applied from t=36. Control clusters receive no intervention thus there will be no change to their population dynamics.

#### The seasonal model

Under this model two alternating seasons, rainy and dry, each 6 generations long, are assumed. The rainy *N_e_* is 30,000, and in the dry season it drops to 10% of the rainy season size. Those in the intervention arm will experience a 90% suppression in both their rainy and dry season *N_e_*’s (Figure 1B).

#### Heterogeneity of clusters

We further explore the effect of heterogeneity among clusters as it is unreasonable to assume all clusters are exactly the same size. The between-cluster variation was introduced by randomly sampling the pre-intervention *N_e_* (or the rainy season *N_e_* under the seasonal model) from a lognormal distribution, with mean *N_e_* equals 30,000 and standard deviation *σ* (in the log scale). Three values of *σ* = (0, 0.1, 0.5) were chosen to represent different magnitudes of heterogeneity. To find *µ* in the log scale we have:

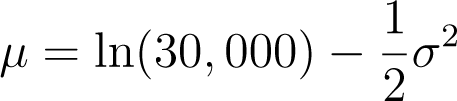

Note that suppression level and seasonality remain as fixed percentages.

### 2.2 Genetic summary statistics

From each simulated dataset, genetic summary statistics were calculated from the 50 sampled individuals from each time point. The following statistics were considered: density of segregating sites, nucleotide diversity (*π*), Tajima’s D and unlinked linkage disequilibrium (LD). The first three were directly obtained via *tskit* (Ralph et al., 2020). For unlinked LD, the *msprime* output was first converted to vcf files as unphased genotypes, then the average from all unlinked *r*^2^ were computed via the Burrows’ method using loci with minor allele frequency of 5% or more (Waples, 2006).

### 2.3 Analysis of cRCT

We aim to study the power of the chosen statistics to differentiate the intervention arm from control under the context of a cRCT and a given scenario. Power was presented as a function of the number of clusters per arm *k* (from 2 to 20) and the time of post-intervention sampling *t*_2_ (from 33 to 0 generations ago). At each iteration, *k* clusters per arm were randomly sampled without replacement from the pools of clusters we previously simulated. The chosen statistic at *t*_2_ were extracted, a two-sided t-test between the arms was performed and the p-value was recorded. The whole process was repeated 1,000 times, and the statistical power was approximated by the proportion of these 1,000 iterations that yielded a statistically significant p-value of *<* 0.05.

To examine the usefulness of incorporating baseline readings, a pre-intervention sample was taken three generations before the intervention at *t*_1_=39. The corresponding t-test was then based on the differences of the summary statistic between *t*_2_ and *t*_1_.

## 3 Results

### 3.1 Constant population model

Statistical powers as a function of the number of clusters (*k*) and the time of sampling (*t*_2_) are presented as heatmaps. Under the homogeneous population scenario (*σ* = 0) without baseline data, all statistics demonstrated distinct detection characteristics for population crashes (left panel of Figure 2). For 99% crashes (results shown in S.I.), LD showed the fastest to equilibrium, but required about 4 to 7 clusters per arm to achieve sufficient power (*>*80%). The density of segregating sites and Tajima’s D achieved high power within 3 generations using 3 clusters per arm, while *π* took 24 generation with the same number of clusters per arm. In the 90% crash scenario (Figure 2), LD and *π* barely showed power exceeding 0.25; other statistics still achieved sufficient power with 3 or 4 clusters per arm, albeit with a longer response time (15 to 21 generations). Overall, Tajima’s D and the density of segregating sites emerged as the most informative statistics for detecting both severe and moderate population crashes, followed by *π* and then LD.

**Figure 2:**
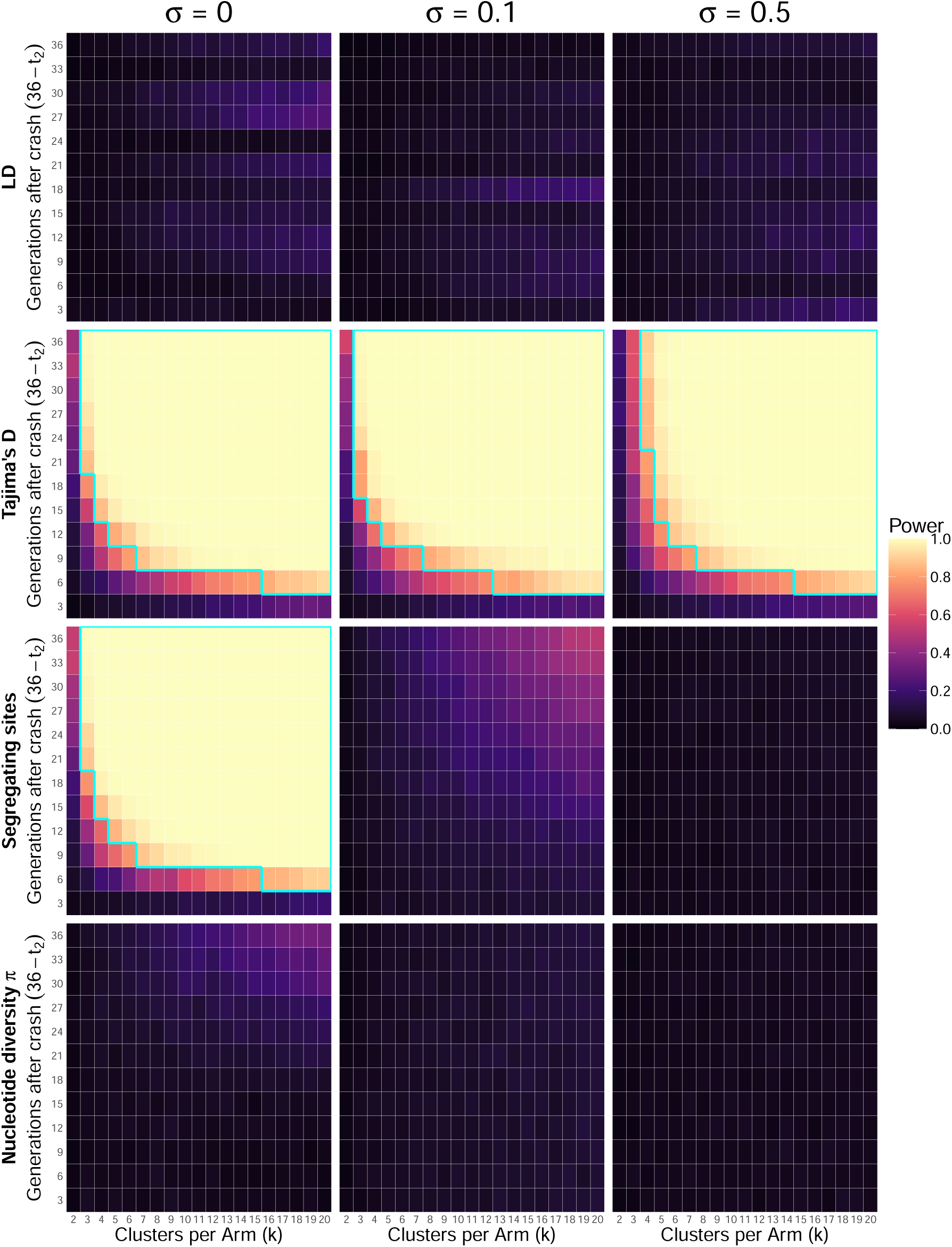
Power of unlinked LD, Tajima’s D, density of segregating sites, and *π* under the constant population model as a function of *k* and *t*_2_, with 90% population suppression. Powers are calculated from cRCT without baseline data, ranged from 0 (darker colours) to 1 (brighter colours). Note that the vertical axis is the number of generations after crash (36 - *t*_2_), with 36 being furthest from the crash. Cyan lines outlines the part where power is greater than 0.8. The three columns display powers under different heterogeneity severities (left: *σ* = 0; middle: *σ* = 0.1; right: *σ* = 0.5).

### 3.2 Seasonal population model

Under seasonal population fluctuations, the result patterns generally paralleled those observed in constant models, though with higher power for equivalent crash severities (Figure 3). An exception was LD, which showed higher power during rainy seasons and power approaching zero during dry seasons. In homogeneous conditions (*σ* = 0) without baseline data, Tajima’s D and density of segregating sites could achieve sufficient detection power within 6 generations with 3 clusters per arm, but *π* required nearly 36 generations and more than 11 clusters. The best choice of statistics remained consistent with constant models, though the seasonal context generally enhanced detection efficiency.

**Figure 3:**
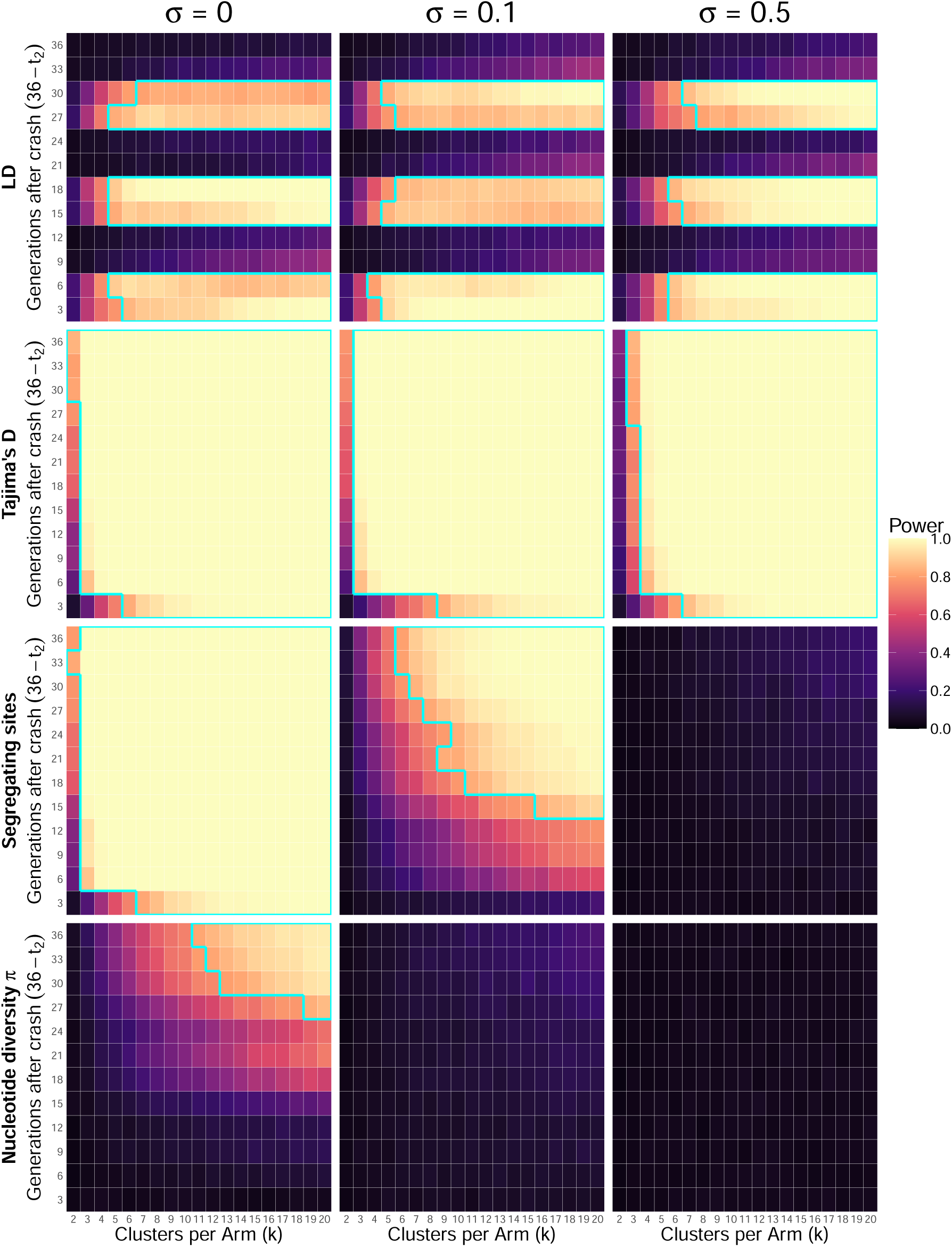
Power of unlinked LD, Tajima’s D, density of segregating sites, and *π* under the seasonal population model as a function of *k* and *t*_2_, with 90% population suppression. Powers are calculated from cRCT without baseline data, ranged from 0 (darker colours) to 1 (brighter colours). Cyan lines outlines the part where power is greater than 0.8. The three columns display powers under different heterogeneity severities (left: *σ* = 0; middle: *σ* = 0.1; right: *σ* = 0.5).

### 3.3 Between-cluster variation

Population heterogeneity (*σ*) had variable effects on power depending on the summary statistic (Figure 2 & 3). Without baseline data, power decreased as the degree of between-cluster variation increased for most statistics, with LD and Tajima’s D being least affected by the variation, maintaining relatively stable power across heterogeneity levels. By contrast, the density of segregating sites and *π* showed substantial power reductions at higher heterogeneity levels. This pattern was consistent across both constant and seasonal population models, suggesting that the impact of heterogeneity on power is largely independent of temporal demographic dynamics.

### 3.4 Baseline collection

The inclusion of baseline data substantially altered the performance of all statistics (results shown in S.I.). Most notably, baseline collection effectively counteracted the negative effects of between-cluster variation, with power remaining relatively constant across *σ* values. The number of segregating sites and *π* benefited most from baseline data, showing significant power increases across all heterogeneity scenarios. In contrast, Tajima’s D derived minimal benefit from baseline collection, while LD showed none or even slightly negative power gains. With baseline data in constant models, all statistics except LD achieved sufficient detection power for 99% crashes within 6 generations using only 2 to 4 clusters per arm. For 90% crashes, Tajima’s D and segregating sites could achieve sufficient power with the same sampling effort but longer response times, while *π* would require at least 4 and better with 5 clusters per arm. In seasonal models, baseline data enabled even faster detection with less clusters.

Overall, baseline data collection altered the optimal choice of statistics. With baseline data, the number of segregating sites was the most robust statistic across both constant and seasonal models, closely followed by Tajima’s D. Without baseline data, Tajima’s D outperformed all other statistics due to its resilience to population heterogeneity.

## 4 Discussion

This work examines how well the summary statistics respond to recent changes in *N_e_*, rather than how they behave as *N_e_* estimators. Our results demonstrated that Tajima’s D followed by density of segregating sites were consistently the best practical choices in terms of statistical power across multiple scenarios. To our surprise, unlinked LD, an estimator previously thought to be useful for early detection of population declines (Antao et al., 2011), did not have significant power especially for small *k*. On paper, its short memory on *N_e_* makes it ideal for quantifying recent demographic changes, but its large variance and coefficient of variation (CV) are the major drawbacks. Suppression only affects the drift component of the observed LD while the sampling portion remains, resulting in small effect size (Waples, 2006). On the other end of the spectrum, *π* is known for its gradual response to demographic changes, containing information of *N_e_* up to hundreds of generations ago (Tajima, 1989), making it unsuitable for cRCT in which the typical trial duration is of 1-3 years. The density of segregating sites has the ability to capture the extinction of rare alleles under strong drift, where *π* = Σ 2*pq* is dominated by highly polymorphic loci. Tajima’s D, being the difference of the two, turned out to be sensitive to recent demographic changes. In our simulations, *D* provided clear signals to population crashes well within the studied time frame of 36 generations post-intervention with high precision.

When *σ* = 0, all clusters have the same pre-intervention condition hence no baseline reading is needed. Power decreases with increasing *σ* as expected when only post-intervention readings at *t*_2_ were used. This effect was particularly pronounced for slow-reacting statistic such as *π*, that powers were mostly below 20 percent across scenarios (Figures 2 & 3). For *σ >* 0, baseline readings are private to each cluster, removing the confounding effect of mixing big and small clusters in the same cRCT. With baseline collection, the differences of the statistic pre- and post-intervention were used, thus the auto-correlation of the statistic plays a key role in determining the standard errors of the t-tests. Taking *π* as an example, *V ar*(*π_t_*_2_ *− π_t_*_1_) = *V ar*(*π_t_*_2_) + *V ar*(*π_t_*_1_) *−* 2*Cov*(*π_t_*_2_ *, π_t_*_1_) has a strong and positive covariance term due to its long memory. The reduction of *V ar*(*π_t_*_2_ *− π_t_*_1_) over the individual variances explains why *π* benefited most from having baseline readings. For fast-reacting statistic like unlinked LD, the auto-correlation is almost zero, and in this case having baseline readings may even be detrimental.

The dry season bottleneck can be viewed as a natural recurrent suppression, which confounds the main intervention effect. Slow-responding statistics were unable to catch up with the switch of seasons hence performed poorly in cRCTs. Interestingly, for unlinked LD, the post-intervention sampling must take place before the end of the rainy season (Fig 3) due to its immediate response. It should note that in some parts of East Africa there are two rainy (and dry) seasons per calendar year, shortening the cycle period. Ideally, the memory-effect of the summary statistic should correlate to the duration of seasonality. On the other hand, the seasonal bottleneck reduces the harmonic mean *N_e_* relative to a constant population of equivalent rainy-season size. Since a smaller *N_e_* amplifies genetic drift, this reduction is expected to increase the divergence in summary statistics between control and intervention arms, which increase the power.

The simulated scenarios and parameters were carefully chosen to resemble the local demography of *An. gambiae s.l.*. The baseline *N_e_* largely follow previous estimates (Hui et al., 2021) from the villages in West Africa, although smaller estimates were obtained from the islands in Lake Victoria, Uganda (Mwima et al., 2026). The choices of magnitude and duration of seasonality were based on previous experiments or modelling (Epopa et al., 2017; Lehmann et al., 2017; Mwima et al., 2024). The ratio of mutation to recombination rate was extracted from existing estimates (Pombi et al., 2006). The simulated genome size (of 2 *×* 1Mb) was much smaller than that of *An. gambiae s.l.* after scaling, thus the statistical power reported here represents a conservative estimate. Higher precision is expected when analysing actual mosquito genomes with far more polymorphic sites. The processing of 50 individuals per cluster per time point was practically achievable and consistent with common field sampling efforts, further supporting the relevance of our power estimates for field studies. Other standard assumptions apply: discrete generations, closed populations, genetically neutral loci.

The powers reported in this work were comparable to those estimated from census collections (Hancock et al., 2025), nonetheless they should not be viewed as a direct comparison because of the different scenarios and assumptions behind the two studies. For fecund species like *Anopheles*, their complex life cycle, high juvenile mortality and high reproductive variance almost guarantee *N_c_ >>> N_e_* (Hedrick, 2005). While census-based methods provide a direct measure of vector density, they are often hindered by high sampling variance and the logistical difficulty of detecting mosquitoes at very low densities. Additionally, *N_c_* is also expected to be noisier and often confounded with the other demographic parameters in the system (Lerch et al., 2026), while *N_e_*, being a composite of these parameters, may be more stable. *N_e_* is a genetic measure for abundance and may lag behind demographic changes, but captures the cumulative effects of a population crash, such as increased drift, inbreeding, and lost in diversity, that snapshot census counts might miss. In principle it may be possible to combine multiple genetic summary statistics to achieve even higher power, potentially via multivariate analysis or statistical learning, subject to further investigation. We can also foresee the possibility of using both *N_c_* and *N_e_* statistics in tandem in the same cRCT, although the improvement will be largely limited by the positive correlation among outcome measures (Vickerstaff et al., 2021).

Interventions with spatial-repelling or dispersal features need to be applied to the entire cluster (e.g. a village) instead of individual households. Spillover of interventions into control clusters could occur through local migration (Multerer et al., 2021). While buffer zones can be introduced between clusters, some genetic summaries are known to be robust against migration, or have a lagged response which could mitigate the spillover effect within the trial period. Beyond entomological endpoints, further assessments are required to correlate vector density suppression to epidemiological endpoints , such as the reduction in malaria incidence or prevalence, in the same or separate trials. There are other practical considerations, for instance, pairing and restricted randomisation of clusters, to further improve the balance and design of cRCTs (Arnold et al., 2024; Hemming & Taljaard, 2023). With new and more effective interventions being constantly developed, the success of field trials also depends on the ability to detect the intervention effect within a reasonable time frame and resources. This work demonstrates a general framework to utilise genetic information in cRCTs, providing a complementary approach to conventional entomological measures and enhancing the power and interpretability of intervention trials.

## 5 Conclusion

This study shows that genetic summary statistics can be effectively used to detect population suppression in clustered randomised controlled trials (cRCTs), but their performance depends on demographic context, between-cluster heterogeneity, and baseline sampling. Across all scenarios without baseline data, Tajima’s D consistently provided the highest and most robust detection power. Unlinked linkage disequilibrium (LD) showed limited practical power due to high variance, despite its rapid response to demographic change. Baseline data collection mitigates the effects of between-cluster heterogeneity and reduces variance, particularly for slow-responding metrics. With baseline data, the density of segregating sites became the most powerful and reliable statistic, closely followed by Tajima’s D, while *π* also showed marked improvements. Seasonal population dynamics generally enhanced detection power, but fast-reacting metrics such as LD were effective only when sampling in rainy seasons. With 50 sampled individuals across all scenarios, cRCT sample size requirements are rather consistent, typically three to five clusters per treatment arm.

## Author contribution

L.Z. coded the simulations. L.Z. & T.H. analysed the data. L.Z. & T.H. wrote the manuscript with inputs from A.B..

## Data and code availability

All codes and simulated data will be available upon publication.

## Acknowledgements

We thank Samantha O’Loughlin, Silke Fuchs, and John Connolly for their useful feedbacks on a previous draft. This work was supported by the Bill & Melinda Gates Foundation and the Open Philanthropy Project, an advised fund of the Silicon Valley Community Foundation. This work is supported by Wellcome Trust (224487).

## Supplementary Information

Constant model, 90% suppression, with baseline

**Figure S1:**
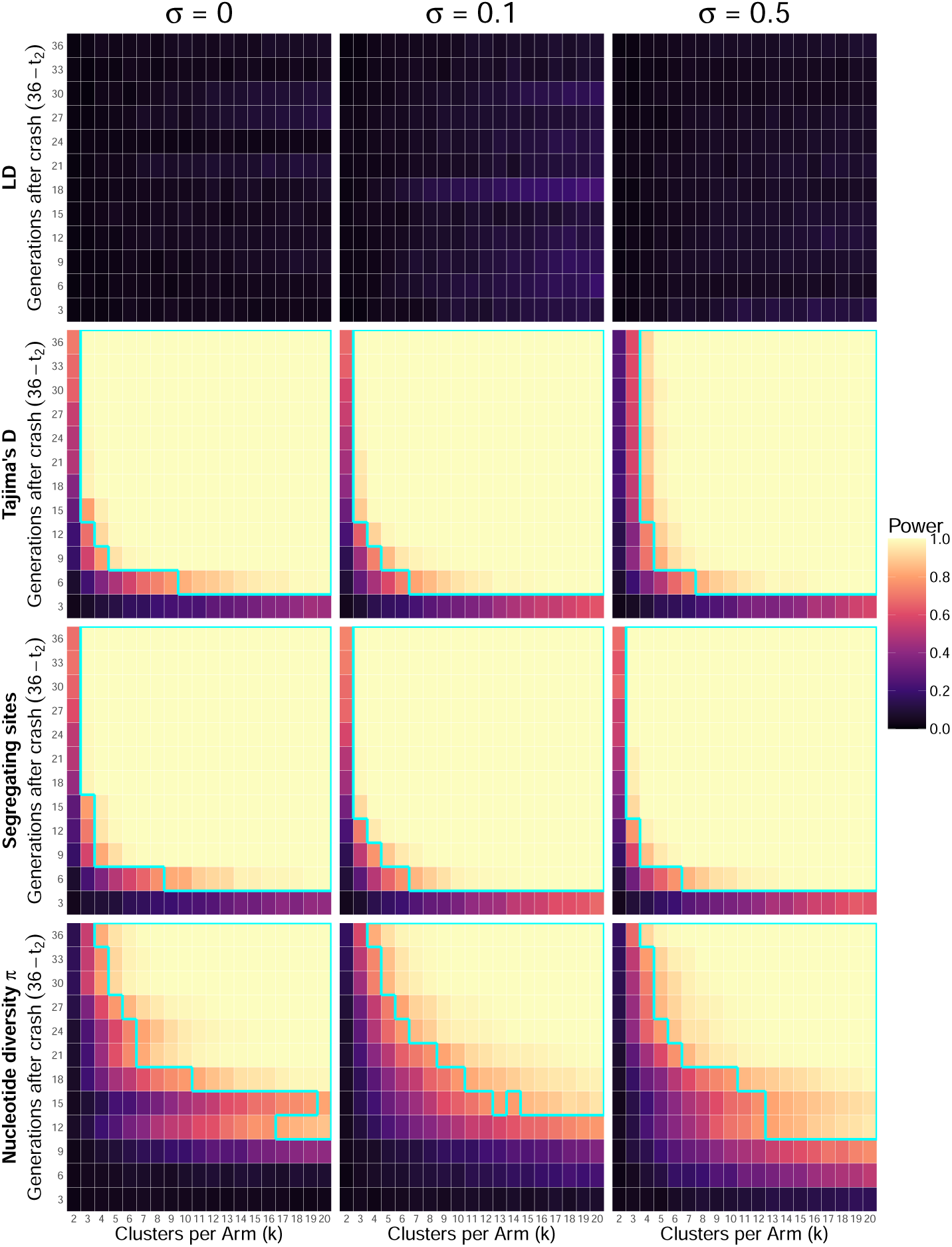
Power of unlinked LD, Tajima’s D, density of segregating sites, and *π* under the constant population model as a function of *k* and *t*_2_, with 90% population suppression. Powers are calculated from cRCT with baseline data, ranged from 0 (darker colours) to 1 (brighter colours). Cyan lines outlines the part where power is greater than 0.8. The three columns display powers under different heterogeneity severities (left: *σ* = 0; middle: *σ* = 0.1; right: *σ* = 0.5).

Seasonal model, 90% suppression, with baseline

**Figure S2:**
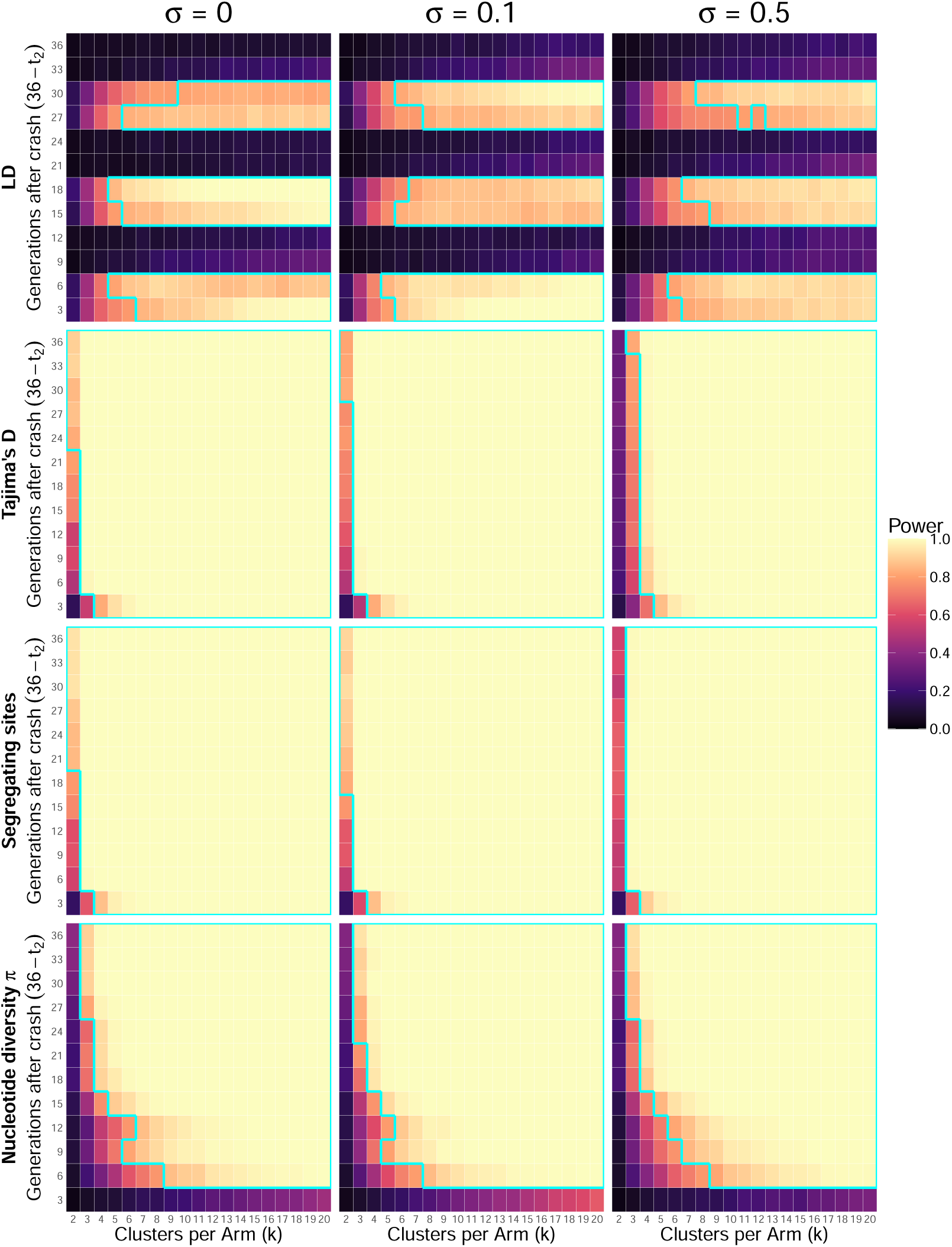
Power of unlinked LD, Tajima’s D, density of segregating sites, and *π* under the seasonal population model as a function of *k* and *t*_2_, with 90% population suppression. Powers are calculated from cRCT with baseline data, ranged from 0 (darker colours) to 1 (brighter colours). Cyan lines outlines the part where power is greater than 0.8. The three columns display powers under different heterogeneity severities (left: *σ* = 0; middle: *σ* = 0.1; right: *σ* = 0.5).

Constant model, 99% suppression, without baseline

**Figure S3:**
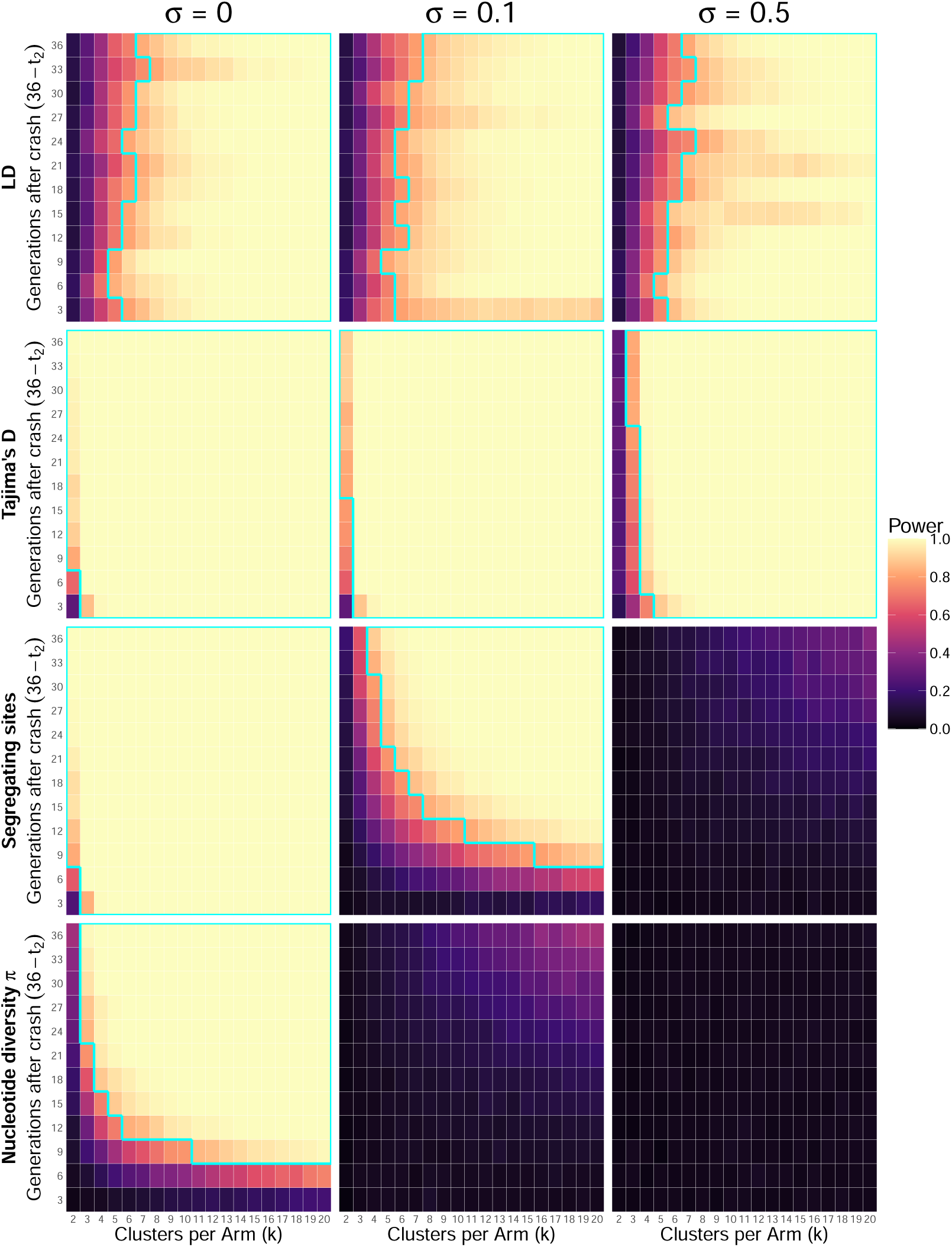
Power of unlinked LD, Tajima’s D, density of segregating sites, and *π* under the constant population model as a function of *k* and *t*_2_, with 99% population suppression. Powers are calculated from cRCT without baseline data, ranged from 0 (darker colours) to 1 (brighter colours). Cyan lines outlines the part where power is greater than 0.8. The three columns display powers under different heterogeneity severities (left: *σ* = 0; middle: *σ* = 0.1; right: *σ* = 0.5).

Constant model, 99% suppression, with baseline

**Figure S4:**
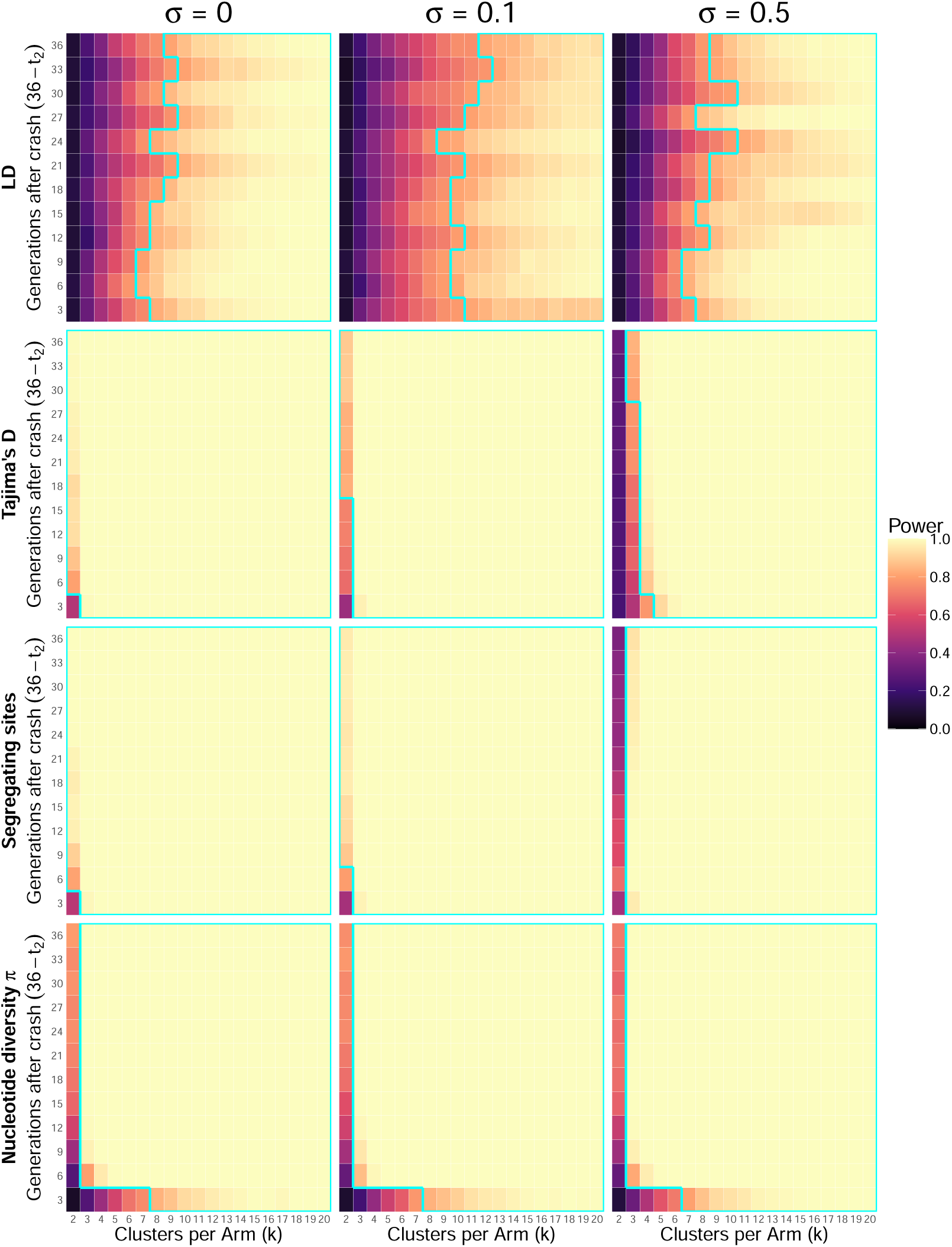
Power of unlinked LD, Tajima’s D, density of segregating sites, and *π* under the constant population model as a function of *k* and *t*_2_, with 99% population suppression. Powers are calculated from cRCT with baseline data, ranged from 0 (darker colours) to 1 (brighter colours). Cyan lines outlines the part where power is greater than 0.8. The three columns display powers under different heterogeneity severities (left: *σ* = 0; middle: *σ* = 0.1; right: *σ* = 0.5).

